# Sarcomere analysis in human cardiomyocytes by computing radial frequency spectra

**DOI:** 10.1101/2025.05.28.655290

**Authors:** Michael Habeck, Nosheen Saleem, Daria Plota, Cleophas Cheruiyot, Tobias Kohl, Stephan Lehnart, Stefan Jakobs, Antje Ebert

## Abstract

In cardiomyocytes, the basic contractile unit are sarcomeres, which are organized in a regular manner facilitating their function. Here, we present a new computational approach to assess the functional properties of sarcomeres at the nanoscale level in human cardiac cells, induced pluripotent stem cell-derived cardiomyocytes (iPSC-CMs). We combined our analysis to different types of high-resolution imaging data, structured illumination microscopy (SIM), stimulated emission depletion (STED) microscopy-based imaging, as well as confocal microscopy data. We show that the radially averaged magnitude spectrum (RAMS) revealed sarcomere properties in a human cardiomyocyte model, iPSC-CMs, and compared our RAMS-based analysis to a real-space approach based on manually selected regions of interest. Moreover, we found the RAMS method suitable to quantify molecular differences of sarcomeres such as present in severe cardiac diseases, such as dilated cardiomyopathy (DCM). Defects in the sarcomere organization that occur in the presence of inherited DCM mutations in sarcomere proteins were efficiently recapitulated by our analysis. This new approach may facilitate streamlined analysis of molecular disease-specific phenotypic imaging data of cardiac cells, aiding our deeper understanding of the molecular basis of cardiac diseases.

## INTRODUCTION

In cardiomyocytes, excitable contractile cells, the smallest contractile unit are the sarcomeres, which generate contractile force in a Ca^2+^-dependent manner by sliding of actin-myosin filaments. Sarcomeres are organized in a highly regular manner, which is critical for their efficient function. Thus, sarcomeres properties are a basic unit of understanding the function of cardiomyocytes. This is critical as sarcomere functions are pathologically altered in a variety of cardiac diseases, including cardiomyopathies, a frequent cause of heart failure. Cardiomyopathies, such as dilated cardiomyopathy, are caused in more than 25-30% of cases by inherited mutations, which typically occur in proteins of the sarcomeres (*1, 2*). Mutations in sarcomere proteins can deform protein structures within the sarcomere, resulting in sarcomere disorganization and altered contractility as well as force generation (*3, 4*). Therefore, accurate quantification of sarcomere properties is a cornerstone not only to structural and computational modelling of cardiomyocyte biomechanics but is vital to molecular studies of human cardiac pathologies. Previous methods of sarcomere analysis typically considered individual imaging strategies (*5-7*). We present here an alternative method for analysis of sarcomere features with multiple imaging technologies and demonstrate its applicability for assessment of disease-specific dysfunctions in sarcomeres. We used a state-of-the-art human cardiac cell model of induced pluripotent stem cell-derived cardiomyocytes (iPSC-CMs). While human iPSC-CMs represent an early stage of heart development (*3, 8*), they are functional human cardiac cells that can be cultured in-vitro and enable to study pathological sarcomere dysfunction in cardiac diseases, such as hypertrophic cardiomyopathy (*9*) and dilated cardiomyopathy (DCM) (*10*).

We employed patient-specific iPSC-CMs derived from an patient with DCM due to an inherited sarcomere protein mutation, tropomyosin (TPM1)-L185F, described earlier (*11, 12*). DCM patient-specific iPSC-CMs were reported to display disrupted sarcomere organization, impaired contractility, and reduced force generation (*7, 11, 13, 14*). Using immunohistochemistry of cardiac markers and confocal-based image analysis, we quantified distinct sarcomere characteristics in DCM iPSC-CMs, compared to WT controls. The new radially averaged magnitude spectrum (RAMS) approach presented here quantifies distances between consecutive z-discs linking adjacent sarcomeres by applying a Fourier-based analysis to establish sarcomere length and sarcomere organization.

We confirmed the RAMS method is applicable to a wider range of state-of-the-art imaging technologies, such as structured illumination microscopy (SIM) and stimulated emission depletion (STED) microscopy-based imaging. Moreover, we corroborated sarcomere analysis with the RAMS method in 3D image layers and compared RAMS to an analysis approach reported earlier. These aspects will enable capturing nanoscale details and facilitate applicability of our approach toward its use with other types of imaging data in the future. Therefore, our findings will assist in gaining deeper insight into the molecular basis associated with DCM in cardiac cells.

## RESULTS

### Computational simulation and Fourier-based analysis of signal intensities

Sarcomeres are defined as the interval or space between two z-discs, to which the actin-based thin filaments attach. Based on this, we applied a new approach to functional analysis of sarcomeres. The z-disc pattern, detected by z-disc-specific proteins such as sarcomeric *α*-actinin, can be considered to correspond to a “wave” that traverses the sarcomere in longitudinal direction. The sarcomere length corresponds to the respective wavelength. Therefore, we applied Fast Fourier transform (FFT) to characterize the wave pattern formed by z-discs without requiring manual processing such as the selection of an ROI or linear segments. We first tested this approach using computational simulation of striated patterns propagating along the *x*-axis (**Fig. 1A**, left) and calculated the respective magnitude spectrum by a 2D FFT (**Fig. 1A**, right). In this idealized simulation, the frequency of the striated pattern was ten, thus the intervals were separated by a distance of 1024/10 = 102.4 pixels. In the Fourier domain, this resulted in distinct peaks at a frequency of *k*_*x*_ = ±10 indicated by red dashed lines (**Fig. 1A**, right). To determine the separating distance, the simulated image intensities (**Fig. 1A**) were projected onto a line (the *x* axis in this example) (**Fig. 1B**). The resulting profile displayed a series of peaks that are separated by a constant distance corresponding to the sarcomere length. The peaks were automatically detected with Otsu thresholding and connected-component labeling (along the *x* axis). The numerical values were extracted via the estimated peak locations (**Fig. 1B**, red dashed lines) and matched the values based on Fourier analysis (**Fig. 1B**). The Fourier approach resulted in an estimated distance of 102.40 pixels, whereas the real-space approach yielded an average value of 102.39 pixels.

**Figure 1:**
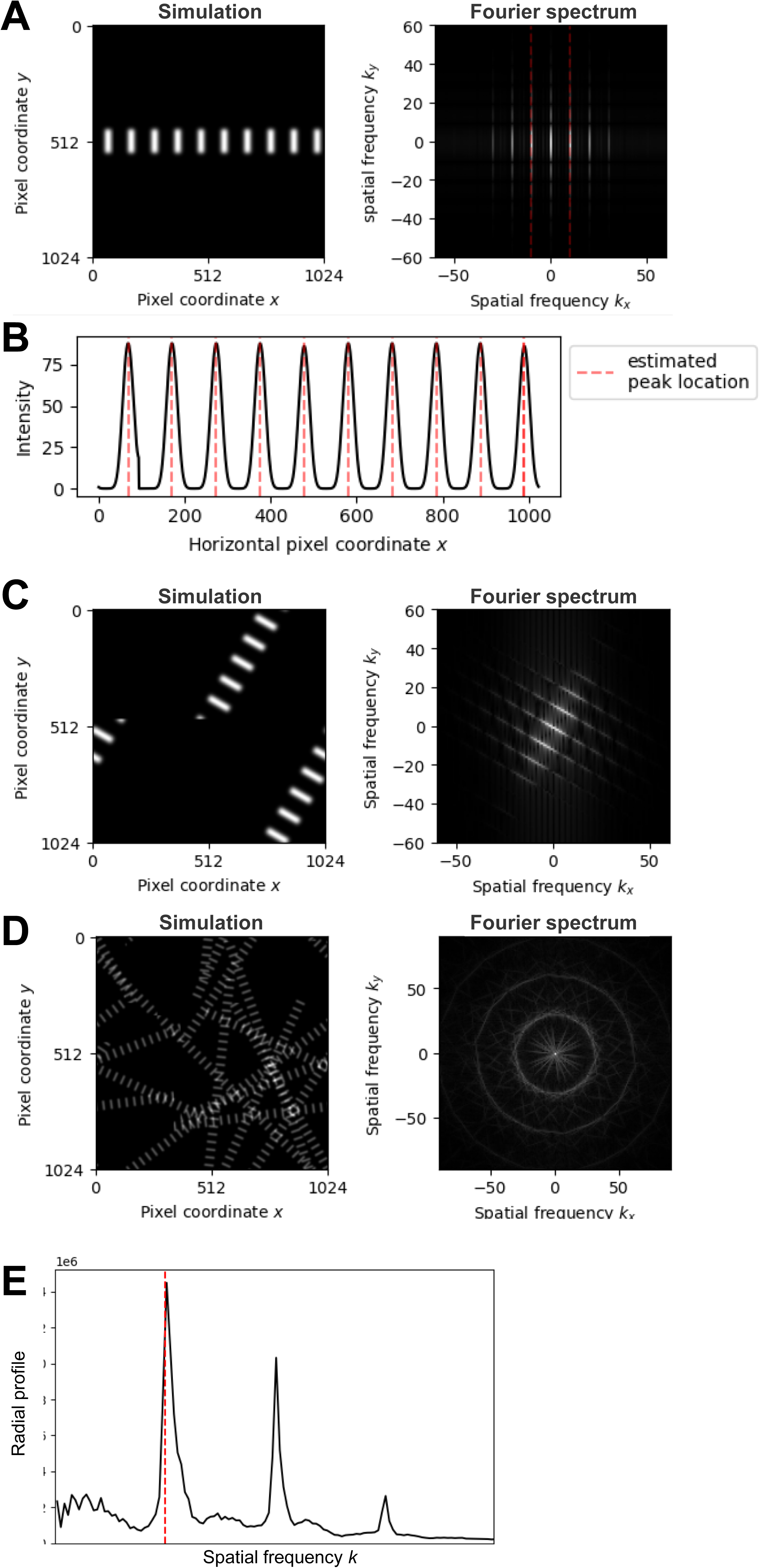
Computational simulation establishes peak maxima extraction via RAMS. **A**, left, computational simulation of striated patterns. **A**, right, magnitude of the 2D Fourier spectrum corresponding to **A**, left. Red dashed lines indicate peaks at a frequency of *k*_*x*_ = ±10, based on the simulation in **A** and corresponding spatial frequencies along the direction of the propagating wave in frequency space. **B**, projection of the simulated image intensities to a one-dimensional profile (*x* axis). Automatically estimated peak locations based on Otsu thresholding are marked by red dashed lines. **C**, left, computational simulation of striated patterns rotated at an angle of 60 degrees. **C**, right, 2D magnitude spectrum corresponding to **C**, left. **D**, left, computational simulation of striated patterns oriented in different spatial directions. **D**, right, magnitude spectrum corresponding to **D**, left. **E**, radially averaged magnitude spectrum (RAMS); red dashed line indicates the global maximum of the RAMS.

In reality, the myofibrils of cardiomyocytes and thus, the striated pattern of sarcomeres can be oriented in many different directions. Therefore, we analyzed the consequences of rotated striation. For example, sarcomeres propagating in vertical direction will have the same Fourier spectrum as shown in Fig. 1A but will be rotated by 90 degrees. Employing computational simulation, we generated striated patterns rotated at an angle of 60 degree (**Fig. 1C**, left), compared to Fig. 1A. Consequently, in the idealized simulation, the respective Fourier spectrum was rotated by the same angle corresponding to its striated pattern (**Fig. 1C**, right).

Sarcomeres present yet differently in early postnatal cardiomyocytes. In this developmentally early stage, immature sarcomeres are oriented in various directions, resulting in a superposition of their Fourier spectra. Assuming isotropic orientation of striated sarcomere patterns in all spatial directions, the Fourier peaks will form a ring around the zero-frequency peak. Thus, we expected to see ring-like structures in the Fourier spectrum of sarcomeres with different orientations as demonstrated by computational simulation (**Fig. 1D**). The radius of the ring corresponded to the dominant frequency or, respectively, the dominant wavelength in the image.

### Computing radial frequency spectra enables extraction of peak maxima

To determine the dominant frequency of striated patterns, we radially averaged the Fourier spectrum. The radially average magnitude spectrum (RAMS) enabled us to estimate the sarcomere length by locating the dominant peak. If *I* denoted the microscopy image and *I*^its 2D Fourier transform with the zero-frequency component shifted to the center, the RAMS is defined as:

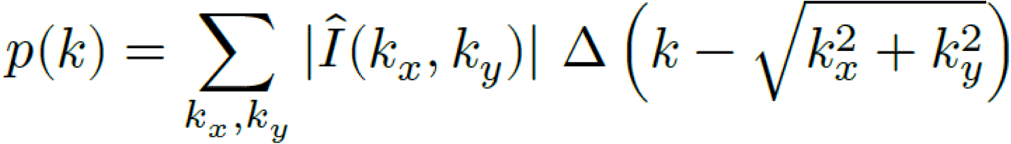

In the definition of the radial profile *p*, the sum runs over all spatial frequencies and the binning function Δ is a rectangular function whose support is determined by a user-defined bin size *δ*:

That is, the radial profile *p* was 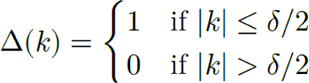 computed by summing up all amplitudes along rings of width *δ* (**Fig. 1D**). Note that in our definition of the radial profile, we omitted normalization by the number of pixels contributing to a ring at frequency *k*, which in two dimensions is proportional to *k*. The first maximum of the radial profile was located at a frequency close to the true frequency of the striated patterns (**Fig. 1E**). With this approach, we estimated the frequency of the striated pattern to be 30.5, which is close to the true value of 30 used in the simulation.

From the estimated frequency *k*_max_, we obtain an estimate *d* of the distance between individual striations:

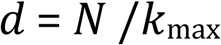

where *N* is the number of pixels. For the simulated pattern, we obtained a distance estimate of 33.57 pixel (the true value is 34.13 pixels).

### Analysis of high-resolution sarcomere imaging of human cardiac tissues by RAMS

Next, we sought to apply the RAMS approach to sarcomere imaging data from human cardiomyocytes. For image analysis of sarcomeres, data generated and reported earlier from human heart tissues from donor hearts rejected for transplantation (*11*) were employed. Following slicing, mounting, fixation and immunostaining for typical cardiac sarcomere markers, cardiac troponin T (TNNT2, TnT) and sarcomeric α-actinin (ACTN2, SAA), confocal imaging was performed as reported earlier (*11*) (**Fig. 2A**). Signals derived from the sarcomere contractile protein, TnT (TNNT2), as well as the z-disc-localized protein SAA (ACTN2), visualized the sarcomeres. As before for computational simulation of striated patterns (**Fig. 1C**), in the sarcomere z-disc patterns detected via SAA (ACTN2) (**Fig. 2B**, left), a regular wave could be considered to traverse the image plane from the lower right to the upper left corner.

**Figure 2:**
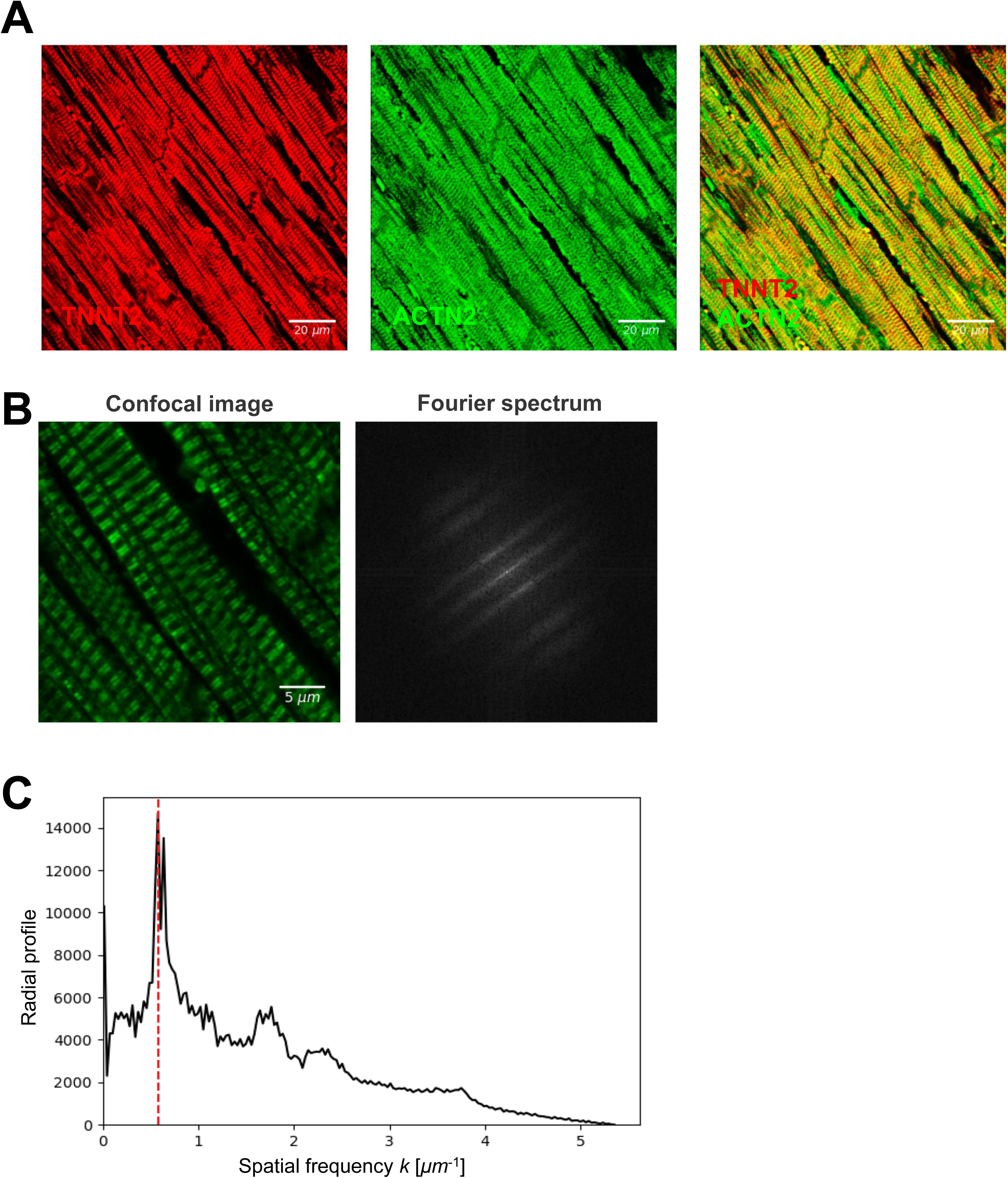
RAMS-based analysis of high-resolution sarcomere imaging of human cardiac tissues. **A**, Human heart tissue section following immunostaining with antibodies against cardiac troponin T (TNNT2), sarcomeric α-actinin (ACTN2). A representative image from an earlier report (*11*) is shown. **B**, left, zoomed-in region of the confocal image shown in **A**, revealing sarcomere z-discs detected by sarcomeric α-actinin (ACTN2); **B**, right, magnitude spectrum obtained with a 2D Fourier transform of the selected region. **C**, radially averaged magnitude spectrum showing peaks at distinct frequencies in Fourier space, based on which the corresponding sarcomere length is computed (red highlighted).

As observed for computational simulation (**Fig 1C**), the Fourier spectrum showed marked peaks close to the central zero-frequency maxima that were oriented longitudinally with the sarcomeres (**Fig 2B**, right). Because the sarcomere filaments in the CMs of adult human heart tissue are aligned in parallel, no ring structure emerged in the magnitude spectrum (**Fig 2B**, right). Based on the Fourier spectrum, the RAMS was obtained (**Fig 2C**). Radial frequency spectra analysis enabled locating the dominant frequency. Accordingly, RAMS allowed us to derive an estimate of the sarcomere length of 1.73 µm.

### Application of RAMS analysis to patient-specific iPSC-CMs

For experimental and translational research, human CMs from adult heart tissues are a most suitable model, however, are difficult to obtain and cannot be cultured nor modified in-vitro (*15*). In this context, human iPSC-CMs, which represent an early post-natal stage of human CMs (*3, 8*), are an alternative model for research purposes. Therefore, human iPSC-CMs were employed to test the RAMS approach for image analysis of sarcomeres. Human iPSC-CMs were derived using previously described protocols (*11*) from the iPSCs of healthy controls (WT) or a DCM patient carrying the inherited DCM mutation TPM1-L185F, both of which were reported earlier (*11*). Human iPSC-CMs derived from these lines and respective protocols were characterized and reported before (*11*). Both WT and DCM patient-specific iPSC-CMs expressed typical cardiac sarcomere markers, including cardiac troponin T (TNNT2, TnT) as well as sarcomeric α-actinin (ACTN2, SAA) (**Fig. 3A**). Human WT and DCM iPSC-CMs were fixed, permeabilized, and subjected to immunostaining via specific antibodies reported earlier (*13*). High-resolution confocal imaging demonstrated striated patterns of WT control or DCM patient-specific iPSC-CMs (**Fig. 3A** and **Fig. 3B**). The confocal image of human WT shown in **Fig. 3A** exhibits striated patterns propagating in multiple spatial directions. As predicted by simulation, the magnitude spectrum shows a strong ring-like pattern (**Fig. 3C**). To determine the radius of the ring, we computed the RAMS by radially averaging the magnitude spectrum, which resulted in a one-dimensional profile with prominent peaks at certain frequencies (**Fig. 3D**). Based on the location of the first peak, the sarcomere length was estimated to be 1.81 µm in WT iPSC-CMs.

**Figure 3:**
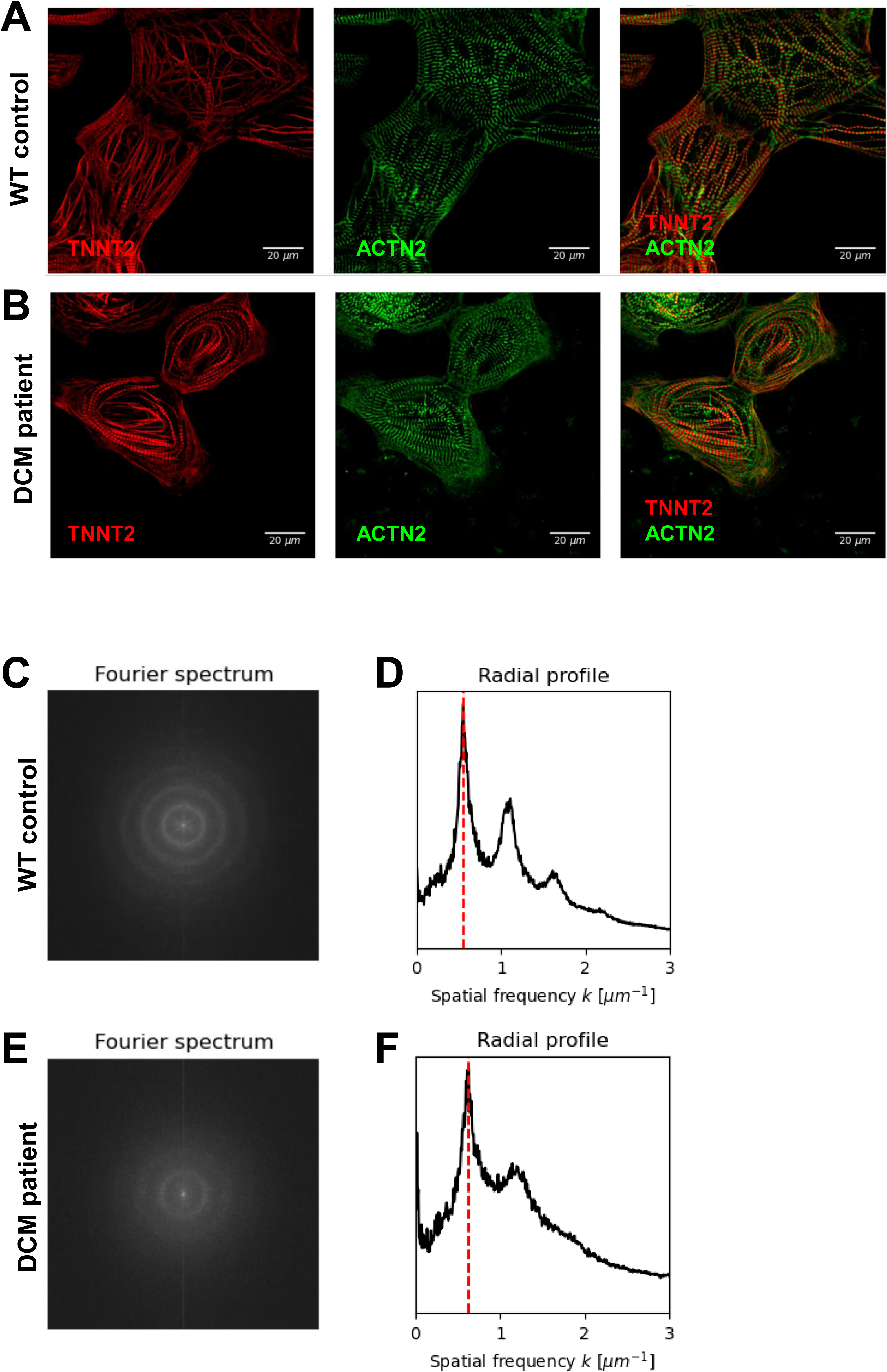
Application of RAMS analysis to sarcomere images from human patient-specific iPSC-CMs. **A**, **B** Representative confocal images of immunostaining for two well-established sarcomere protein markers, cardiac troponin T (TNNT2, TnT), and sarcomeric alpha-actinin (ACTIN2, SAA) in WT (control) iPSC-CMs (**A**) compared to DCM patient-specific iPSC-CMs carrying a disease-causing mutation in tropomyosin, TPM1-L185F (**B**). The images are part of an earlier report (*11*). **C**, **F**, magnitude spectra obtained with a 2D Fourier transform of the images shown in **A** and **B**, respectively; **D**, **G**, radial profiles for the Fourier spectra shown in **C**, **F**; red dashed line indicates the dominant frequency for the respective images. **A, C, D**, WT (control) iPSC-CMs; **B, E, G**, DCM patient-specific iPSC-CMs.

Next, we applied the RAMS-based analysis also to high-resolution images of DCM patient-specific iPSC-CMs (**Fig. 3B**). Again, a ring pattern emerged in the Fourier domain (**Fig. 3E**) albeit weaker than for WT iPSC-CMs suggesting the sarcomere structure is less well organized in DCM iPSC-CMs.

Using the RAMS (**Fig. 3F**) the sarcomere length was found to be reduced, in line with our previous publication of the respective image data set (*11*). The sarcomere length was estimated to be 1.60 µm for DCM iPSC-CMs. Due to the inherited mutation of a contractile protein, TPM1, the DCM patient-specific iPSC-CMs displayed a lower degree of organization, which was reflected in the diffuse ring pattern and wider peak ranges of the respective radial profile (**Fig. 3F**).

### Comparison of RAMS to segmentation-based signal processing from imaging data

To validate our approach, we compared RAMS-based sarcomere analysis to a different method previously reported (*13, 14, 16*). To establish sarcomere properties, the reference method applied analysis based on region of interests (ROIs). As successive z-disc separate and link adjacent sarcomeres, measuring the distance between two successive z-lines allowed us to estimate the length of the sarcomere with its border marked by the two z-discs. To estimate the length of sarcomeres using high-resolution confocal microscopy images of iPSC-CMs (WT control) (**Fig. 4A**), we utilized intensity projections along each linear segment and thereby obtained a one-dimensional curve that exhibits local maxima (**Fig. 4B**). Following application of Otsu thresholding, the distances between successive peaks were computed and enabled us to estimate the sarcomere length (**Fig. 4C-D**). Using all ROIs, we obtained 81 estimates of the sarcomere length in WT iPSC-CMs with a mean and standard deviation of 1.84 ± 0.14 µm. In comparison, the RAMS-based method delivered a value (1.81 µm) in good agreement with the ROI-based analysis.

**Figure 4:**
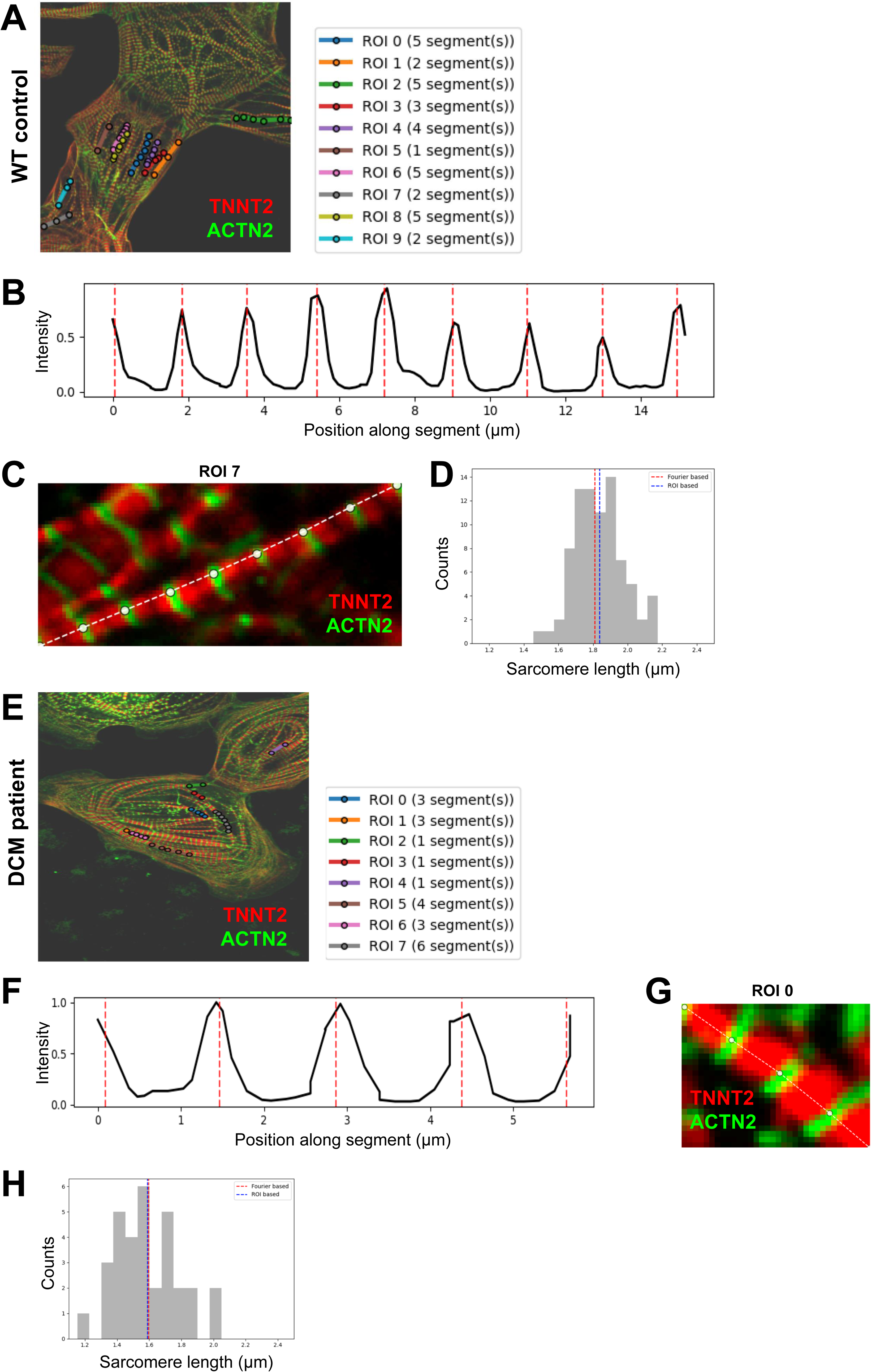
Comparison of RAMS to segmentation-based signal processing from imaging data to establish sarcomere properties. **A**, Segmentation via regions of interest (ROIs) to collect TNNT2 and SAA-dependent signals for WT iPSC-CM sarcomeres based on intensity projection along linear segments. A representative image, which is part of an earlier report (*11*), is shown. **B**, One-dimensional curve exhibiting local maxima based on intensity projections. **C**, Computed peak locations after Otsu thresholding of the intensity projection. **D**, Estimation of the sarcomere length based on ROI analysis (gray histogram, blue dashed line indicates the mean value) or RAMS analysis (red dashed line). **E-H**, analysis corresponding to **A-D** for DCM patient-specific iPSC-CM sarcomeres.

When applying the same procedure to confocal images of DCM patient-specific iPSC-CMs, we obtained results (**Fig. 4E-H**) in line with the above findings. From all ROIs, we obtained 32 measurements of the sarcomere length with a mean value and standard deviation of 1.59 ± 0.20 µm. Again, the value obtained with the RAMS analysis (1.60 µm) were in good agreement with the ROI-based analysis.

### Application of RAMS to imaging data from different high-resolution microscopy techniques

We next explored whether the RAMS-based analysis would be qualified to analyze sarcomere parameters of other high-resolution imaging technologies. We selected Structured Illumination Microscopy (SIM) (**Fig. 5A**) as well as Stimulated Emission Depletion (STED) microscopy images of DCM iPSC-CMs (**Fig. 5B**). Application of spectral analysis to SIM (**Fig. 5C**) and STED (**Fig. 5F**) data enabled us to compute magnitude spectra via 2D FFT (**Fig. 5D, G**), based on which we computed radially averaged frequency spectra (**Fig. 5E, H**) and derived the sarcomere length. The estimated sarcomere length for the SIM image was 1.58 µm. Analysis of the STED image gave a similar value of 1.66 µm.

**Figure 5:**
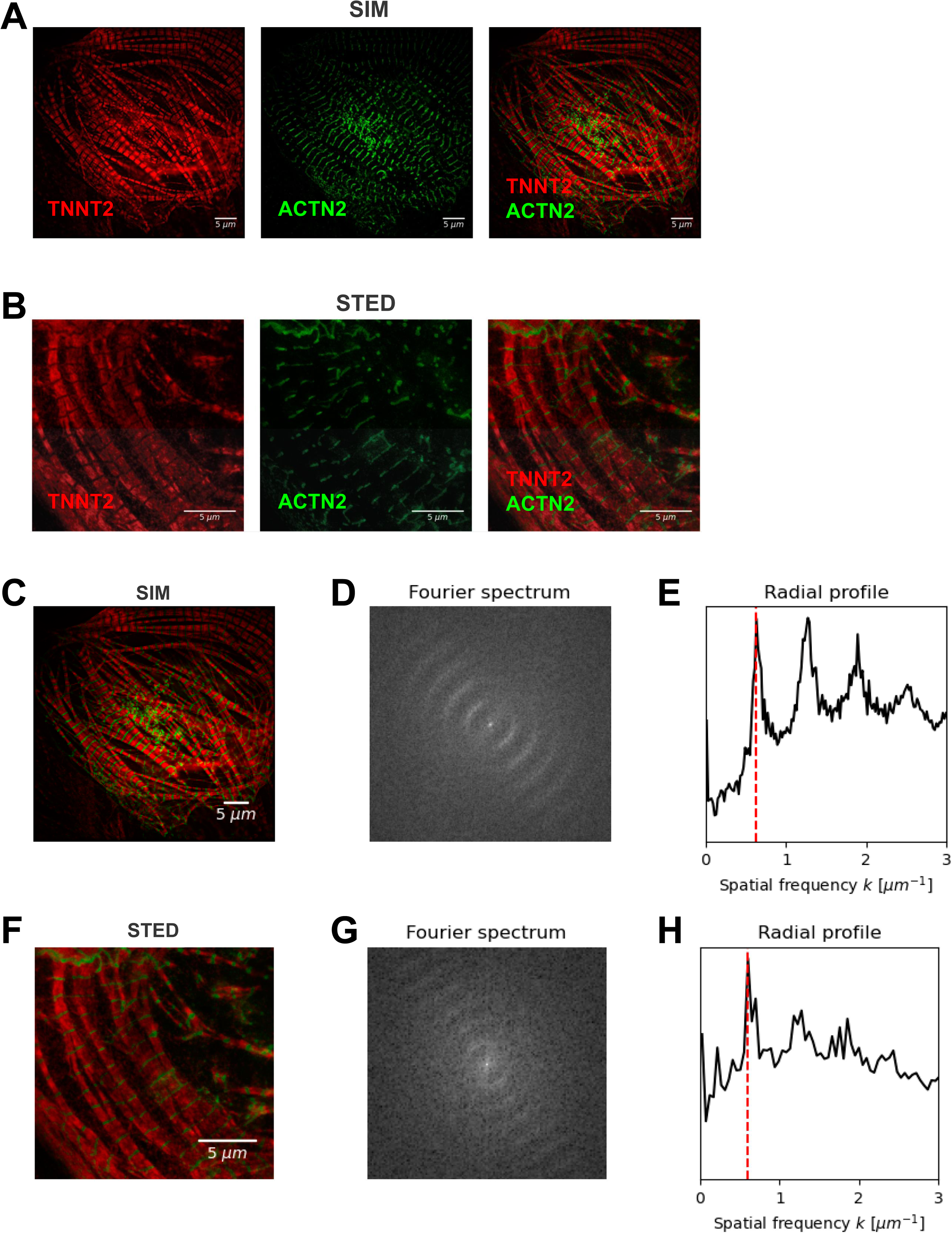
RAMS-based analysis of different high-resolution microscopy imaging data. **A**, Structured Illumination Microscopy (SIM) and **B**, Stimulated Emission Depletion (STED) microscopy images of DCM patient-specific iPSC-CMs previously subjected to immunostaining for cardiac troponin T (TNNT2, TnT), and sarcomeric alpha-actinin (ACTN2, SAA) as described in Fig. 3. **C**, SIM and **F**, STED images were used to compute magnitude spectra (**D**, **G**). (**E**, **H**) Analysis of radially averaged magnitude spectra. Red dashed line, dominant frequency or wavelength for the respective imaging data.

Together, the RAMS-based method enabled us to analyze sarcomere properties such as sarcomere length and organization in high-resolution imaging data derived from human patient-specific iPSC-CMs as well as controls. Moreover, RAMS analysis can be applied to data obtained via different high-resolution imaging techniques.

## DISCUSSION

Sarcomeres are the basic units generating contractile force in striated muscle. Their regularity is crucial for regular function of muscle cells and is altered in severe cardiac diseases such as cardiomyopathy and heart failure (*3, 4*). Several previous reports presented strategies for sarcomere measurements, which retrieved the sarcomere length as a measure of z-disc distances and by defining a point of reference followed by Fast Fourier transform (*5-7*). Moreover, based on manual selection, intensity profiles and feature extraction were combined to conveniently extract multiple as well as individual sarcomere intervals (*17*). Other studies focused on live cell analysis via movie recording and employed image segmentation to derive parameters corresponding to the entire visualized population of sarcomeres (*18*) including computation of wavelet peaks (*19*). In contrast, the radially average magnitude spectrum (RAMS) approach characterizes functional sarcomere properties such as sarcomere length and organization in an automated manner using entire images without the need for manual selection. In addition, radial averaging provides not only the mean sarcomere length of the respective imaging data, but also a profile of Fourier space frequencie*s* thereby offering insights into the regularity of sarcomere organization. The RAMS-based method enables us to analyze the sarcomere properties of human CMs from different sources, such as adult human cardiac tissue biopsies as well as human iPSC-CMs. Moreover, RAMS analysis can be applied to data obtained via different high-resolution imaging techniques, such as confocal and structured illumination microscopy (SIM) as well as simulated emission depletion (STED) imaging. Thus, we present a new method that allows for an analysis of cardiomyocyte sarcomeres of different source and developmental stage – iPSC-derived CMs, adult CMs – with various high-resolution imaging strategies. We also show the RAMS approach to work reliably for the characterization of sarcomere properties from patients with severe cardiac disease, such as DCM due to inherited mutations.

We denoted the applicability of the RAMS approach also by validating the results compared to an earlier method for sarcomere analysis which uses manual feature selection (*13*). This demonstrates the scope of techniques for which the RAMS method can be employed but also its potential for analysis of different types of data derived with other imaging approaches. In the future, the scope of the RAMS method could be extended by integrating machine learning or deep learning strategies for image segmentation. Together, we consider our approach a new analysis method and starting point for further quantitative modelling of sarcomere parameters. Thus, our method could support and be applied in future studies of functional cardiomyocyte biomechanics as well as in translational cardiac biology investigating severe cardiac diseases such as DCM.

## MATERIALS AND METHODS

### Generation, culture, and cardiac differentiation of human iPSCs

All protocols for studies with human iPSC were approved by the Goettingen University Ethical Board (15/2/20 and 20/9/16An) and the Odense University Ethical Board (Projekt ID S-20140073HLP). Informed consent was obtained from all participants and all research was performed in accordance with relevant guidelines and regulations. WT iPSCs were derived and characterized as described before (*11*). DCM patient-specific iPSCs carrying the DCM mutation tropomyosin (TPM1) L185F were derived and characterized before (*11*). Human iPSCs were cultured on Matrigel-coated plates (Corning) utilizing chemically defined E8 medium as reported previously. Human iPSCs were passaged every four days via EDTA (Life Technologies) (*20, 21*), and the culture medium was changed daily. A small molecule-based approach was used to generate beating cardiomyocytes from human iPSCs, as reported previously (*13, 22-25*). The resulting human iPSC-CMs were subsequently cultured in RPMI (Life Technologies) with B27 supplement (Life Technologies). On day 20-25 of cardiac differentiation, the beating iPSC-CMs were dissociated using trypleE (Life Technologies), and used for experiments.

### Immunostaining and confocal imaging

The human iPSC-CMs were plated on coverslips and allowed to recover for 2 to 3 days. Cells were fixed in 4% paraformaldehyde and stained with primary antibodies against cardiac troponin T (TNNT2, TnT, rabbit; Thermo Fisher Scientific) and sarcomeric alpha-actinin (ACTIN2, SAA, mouse, Sigma-Aldrich) overnight. Anti-Mouse IgG1 Alexa Fluor 488 and anti-Rabbit IgG (H+L; heavy light chain) Alexa Fluor 568 were used as secondary antibodies to stain coverslips. Coverslips were mounted onto glass slides and imaging was performed using a confocal microscope (LSM 710 Meta, Zeiss) equipped with a 63X oil-immersed objective, as described previously (*13, 16, 26*)

### Sarcomere Length Analysis

ROI-based image analysis was performed by ImageJ according to previously described methods (*13, 14, 16*). From the line segments, projected intensity profiles were computed with an in-house Python routine. Peaks in the 1D real-space profiles were detected and located with Otsu thresholding as offered by Sci-kit image and connected-component labeling. Analysis via the radially averaged magnitude spectrum (RAMS) method was performed on images of the sarcomere z-disc-localized protein SAA (ACTN2) which were subjected to Fourier analysis. Based on the magnitude spectra, radial profiles in frequency space were generated and the dominant frequency was derived from the peak location. The analysis was implemented and performed in Python version 3.12 and built on the Python packages NumPy, SciPy, Scikit-image and Matplotlib.

### Stimulated emission depletion microscopy (STED)

Human iPSC-CMs were plated on coverslips between day 22 to 26 of cardiac differentiation. Following fixation, permeabilization, and blocking, and primary antibody staining as described above, secondary antibodies (goat anti-rabbit STAR635P, ST635P-1002-500UG, Abberior, and goat anti-mouse STAR580, ST580-1001-500UG, Abberior) were applied. Finally, coverslips were mounted in ProLong Gold Antifade Mountant (Thermo Fisher Scientific). Confocal and STED images were acquired using a Leica TCS SP8 system with a HC PL APO C2S 100x/1.40 oil objective. Dual color STED was performed according to the spectral properties of fluorophores STAR 580 and STAR 635P, with excitation at 580nm and 635nm wavelength by a white-light laser and detection between 590-625 nm and 646-699 nm, respectively. Scanning was performed at 20nm x20nm pixel size, pixel dwell time 480 ns, scanning speed 600 Hz, 12to 16-fold line accumulation, and STED depletion at 775 nm. Raw images were processed in Fiji ImageJ.

### Structured Illumination Microscopy (SIM)

Human iPSC-CMs were harvested between day 22 to 26 of cardiac differentiation and plated on coverslips, followed by fixation and immunostaining as above. 2D-SIM data were recorded using a MI-SIM (CSR Biotech Co., Ltd. (Guangzhou, China) microscope equipped with a 100x/1.5NA oil objective (UPLAPO100XOHR) and an ORCA-Fusion BT (Hamamatsu) sCMOS camera essentially as described previously (*27*). Images were acquired in the 2D-SIM-3 mode. Raw data were processed by Wiener reconstruction followed by sparse deconvolution (*28*).

## Supporting information

Habeck et al_supplemental figure

## ACKNOWLEDGEMENTS

This study was supported by the Deutsche Forschungsgemeinschaft (German Research Foundation), Sonderforschungsbereich 1002, project A12 (A.E.); Deutsche Forschungsgemeinschaft, Grant/Award Number: RTG 2824; and under Germany’s Excellence Strategy - EXC 2067/1- 390729940. D.P. is supported as associated doctoral student of the RTG2824 funded by the Deutsche Forschungsgemeinschaft. C.C. is supported as doctoral student of the RTG2824 funded by the Deutsche Forschungsgemeinschaft. We are grateful for support by the DZHK (German Center for Cardiovascular Research), partner site Goettingen, Germany, project IDs 81X4300123, 81X1300131. We thank for funding by the Clinic for Cardiology and Pneumology at the University Medical Center, Goettingen University and support by the Central Service Unit for Cell Sorting at the University Medical Center, Goettingen University. M. H. gratefully acknowledges funding by the Carl Zeiss Foundation within the program “CZS Stiftungsprofessuren”. We thank CSR Biotech Co., Ltd. (Guangzhou, China) for granting generous access to a live-cell MI-SIM super-resolution microscope, and help with data acquisition, image reconstruction, analysis as well as discussions.

---

**SUPPLEMENTARY FIGURE 1**

**STED after histogram equalization**

The STED image described and shown in Fig. 5B was subjected to histogram equalization.

## REFERENCES

1. R. Eldemire, L. Mestroni, M. R. G. Taylor, Genetics of Dilated Cardiomyopathy. Annu Rev Med 75, 417–426 (2024).

2. E. M. McNally, L. Mestroni, Dilated Cardiomyopathy: Genetic Determinants and Mechanisms. Circ Res 121, 731–748 (2017).

3. T. Eschenhagen, L. Carrier, Cardiomyopathy phenotypes in human-induced pluripotent stem cell-derived cardiomyocytes-a systematic review. Pflugers Arch 471, 755–768 (2019).

4. J. van der Velden, G. J. M. Stienen, Cardiac Disorders and Pathophysiology of Sarcomeric Proteins. Physiol Rev 99, 381–426 (2019).

5. C. M. Douglas, J. E. Bird, D. Kopinke, K. A. Esser, An optimized approach to study nanoscale sarcomere structure utilizing super-resolution microscopy with nanobodies. PLoS One 19, e0300348 (2024).

6. A. Chopra, M. L. Kutys, K. Zhang, W. J. Polacheck, C. C. Sheng, R. J. Luu, J. Eyckmans, J. T. Hinson, J. G. Seidman, C. E. Seidman, C. S. Chen, Force Generation via beta-Cardiac Myosin, Titin, and alpha-Actinin Drives Cardiac Sarcomere Assembly from Cell-Matrix Adhesions. Dev Cell 44, 87–96 e85 (2018).

7. J. T. Hinson, A. Chopra, N. Nafissi, W. J. Polacheck, C. C. Benson, S. Swist, J. Gorham, L. Yang, S. Schafer, C. C. Sheng, A. Haghighi, J. Homsy, N. Hubner, G. Church, S. A. Cook, W. A. Linke, C. S. Chen, J. G. Seidman, C. E. Seidman, HEART DISEASE. Titin mutations in iPS cells define sarcomere insufficiency as a cause of dilated cardiomyopathy. Science 349, 982–986 (2015).

8. A. D. Ebert, S. Diecke, I. Y. Chen, J. C. Wu, Reprogramming and transdifferentiation for cardiovascular development and regenerative medicine: where do we stand? EMBO Mol Med 7, 1090–1103 (2015).

9. F. Lan, A. S. Lee, P. Liang, V. Sanchez-Freire, P. K. Nguyen, L. Wang, L. Han, M. Yen, Y. Wang, N. Sun, O. J. Abilez, S. Hu, A. D. Ebert, E. G. Navarrete, C. S. Simmons, M. Wheeler, B. Pruitt, R. Lewis, Y. Yamaguchi, E. A. Ashley, D. M. Bers, R. C. Robbins, M. T. Longaker, J. C. Wu, Abnormal calcium handling properties underlie familial hypertrophic cardiomyopathy pathology in patient-specific induced pluripotent stem cells. Cell Stem Cell 12, 101–113 (2013).

10. N. Sun, M. Yazawa, J. Liu, L. Han, V. Sanchez-Freire, O. J. Abilez, E. G. Navarrete, S. Hu, L. Wang, A. Lee, A. Pavlovic, S. Lin, R. Chen, R. J. Hajjar, M. P. Snyder, R. E. Dolmetsch, M. J. Butte, E. A. Ashley, M. T. Longaker, R. C. Robbins, J. C. Wu, Patient-specific induced pluripotent stem cells as a model for familial dilated cardiomyopathy. Sci Transl Med 4, 130ra147 (2012).

11. Y. Dai, N. Ignatyeva, H. Xu, R. Wali, K. Toischer, S. Brandenburg, C. Lenz, J. Pronto, F. E. Fakuade, S. Sossalla, E. M. Zeisberg, A. Janshoff, I. Kutschka, N. Voigt, H. Urlaub, T. B. Rasmussen, J. Mogensen, S. E. Lehnart, G. Hasenfuss, A. Ebert, An Alternative Mechanism of Subcellular Iron Uptake Deficiency in Cardiomyocytes. Circ Res 133, e19–e46 (2023).

12. H. N. Saleem, N. Ignatyeva, C. Stuut, S. Jakobs, M. Habeck, A. Ebert, 3D Computational Modeling of Defective Early Endosome Distribution in Human iPSC-Based Cardiomyopathy Models. Cells 13, (2024).

13. Y. Dai, A. Amenov, N. Ignatyeva, A. Koschinski, H. Xu, P. L. Soong, M. Tiburcy, W. A. Linke, M. Zaccolo, G. Hasenfuss, W. H. Zimmermann, A. Ebert, Troponin destabilization impairs sarcomere-cytoskeleton interactions in iPSC-derived cardiomyocytes from dilated cardiomyopathy patients. Sci Rep 10, 209 (2020).

14. A. V. Malkovskiy, N. Ignatyeva, Y. Dai, G. Hasenfuss, J. Rajadas, A. Ebert, Integrated Ca2+ flux and AFM force analysis in human iPSC-derived cardiomyocytes. Biol Chem, (2020).

15. M. T. Zhao, S. Ye, J. Su, V. Garg, Cardiomyocyte Proliferation and Maturation: Two Sides of the Same Coin for Heart Regeneration. Front Cell Dev Biol 8, 594226 (2020).

16. H. Xu, R. Wali, C. Cheruiyot, J. Bodenschatz, G. Hasenfuss, A. Janshoff, M. Habeck, A. Ebert, Non-negative blind deconvolution for signal processing in a CRISPR-edited iPSC-cardiomyocyte model of dilated cardiomyopathy. FEBS Lett, (2021).

17. M. Baheux Blin, V. Loreau, F. Schnorrer, P. Mangeol, PatternJ: an ImageJ toolset for the automated and quantitative analysis of regular spatial patterns found in sarcomeres, axons, somites, and more. Biol Open 13, (2024).

18. B. Zhao, K. Zhang, C. S. Chen, E. Lejeune, Sarc-Graph: Automated segmentation, tracking, and analysis of sarcomeres in hiPSC-derived cardiomyocytes. PLoS Comput Biol 17, e1009443 (2021).

19. C. N. Toepfer, A. Sharma, M. Cicconet, A. C. Garfinkel, M. Mucke, M. Neyazi, J. A. L. Willcox, R. Agarwal, M. Schmid, J. Rao, J. Ewoldt, O. Pourquie, A. Chopra, C. S. Chen, J. G. Seidman, C. E. Seidman, SarcTrack. Circ Res 124, 1172–1183 (2019).

20. G. Chen, D. R. Gulbranson, Z. Hou, J. M. Bolin, V. Ruotti, M. D. Probasco, K. Smuga-Otto, S. E. Howden, N. R. Diol, N. E. Propson, R. Wagner, G. O. Lee, J. Antosiewicz-Bourget, J. M. Teng, J. A. Thomson, Chemically defined conditions for human iPSC derivation and culture. Nat Methods 8, 424–429 (2011).

21. A. D. Ebert, K. Kodo, P. Liang, H. Wu, B. C. Huber, J. Riegler, J. Churko, J. Lee, P. de Almeida, F. Lan, S. Diecke, P. W. Burridge, J. D. Gold, D. Mochly-Rosen, J. C. Wu, Characterization of the molecular mechanisms underlying increased ischemic damage in the aldehyde dehydrogenase 2 genetic polymorphism using a human induced pluripotent stem cell model system. Sci Transl Med 6, 255ra130 (2014).

22. X. Lian, J. Zhang, S. M. Azarin, K. Zhu, L. B. Hazeltine, X. Bao, C. Hsiao, T. J. Kamp, S. P. Palecek, Directed cardiomyocyte differentiation from human pluripotent stem cells by modulating Wnt/beta-catenin signaling under fully defined conditions. Nat Protoc 8, 162–175 (2013).

23. X. Lian, C. Hsiao, G. Wilson, K. Zhu, L. B. Hazeltine, S. M. Azarin, K. K. Raval, J. Zhang, T. J. Kamp, S. P. Palecek, Robust cardiomyocyte differentiation from human pluripotent stem cells via temporal modulation of canonical Wnt signaling. Proc Natl Acad Sci U S A 109, E1848–1857 (2012).

24. A. Ebert, A. U. Joshi, S. Andorf, Y. Dai, S. Sampathkumar, H. Chen, Y. Li, P. Garg, K. Toischer, G. Hasenfuss, D. Mochly-Rosen, J. C. Wu, Proteasome-Dependent Regulation of Distinct Metabolic States During Long-Term Culture of Human iPSC-Derived Cardiomyocytes. Circ Res 125, 90–103 (2019).

25. Y. J. Shu, J. F. He, R. J. Pei, P. He, Z. H. Huang, S. M. Chen, Z. Q. Ou, J. L. Deng, P. Y. Zeng, J. Zhou, Y. Q. Min, F. Deng, H. Peng, Z. Zhang, B. Wang, Z. H. Xu, W. X. Guan, Z. Y. Hu, J. K. Zhang, Immunogenicity and safety of a recombinant fusion protein vaccine (V-01) against coronavirus disease 2019 in healthy adults: a randomized, double-blind, placebo-controlled, phase II trial. Chin Med J (Engl), (2021).

26. A. V. Malkovskiy, N. Ignatyeva, Y. Dai, G. Hasenfuss, J. Rajadas, A. Ebert, Integrated Ca(2+) flux and AFM force analysis in human iPSC-derived cardiomyocytes. Biol Chem 402, 113–121 (2020).

27. X. Huang, J. Fan, L. Li, H. Liu, R. Wu, Y. Wu, L. Wei, H. Mao, A. Lal, P. Xi, L. Tang, Y. Zhang, Y. Liu, S. Tan, L. Chen, Fast, long-term, super-resolution imaging with Hessian structured illumination microscopy. Nat Biotechnol 36, 451–459 (2018).

28. W. Zhao, S. Zhao, L. Li, X. Huang, S. Xing, Y. Zhang, G. Qiu, Z. Han, Y. Shang, D. E. Sun, C. Shan, R. Wu, L. Gu, S. Zhang, R. Chen, J. Xiao, Y. Mo, J. Wang, W. Ji, X. Chen, B. Ding, Y. Liu, H. Mao, B. L. Song, J. Tan, J. Liu, H. Li, L. Chen, Sparse deconvolution improves the resolution of live-cell super-resolution fluorescence microscopy. Nat Biotechnol 40, 606–617 (2022).

