## Supplementary figures and images for "Sarcomere analysis in human cardiomyocytes by computing radial frequency spectra"

### Habeck et al_supplemental figure

**Supplemental Figure 1**

STED after histogram equalization

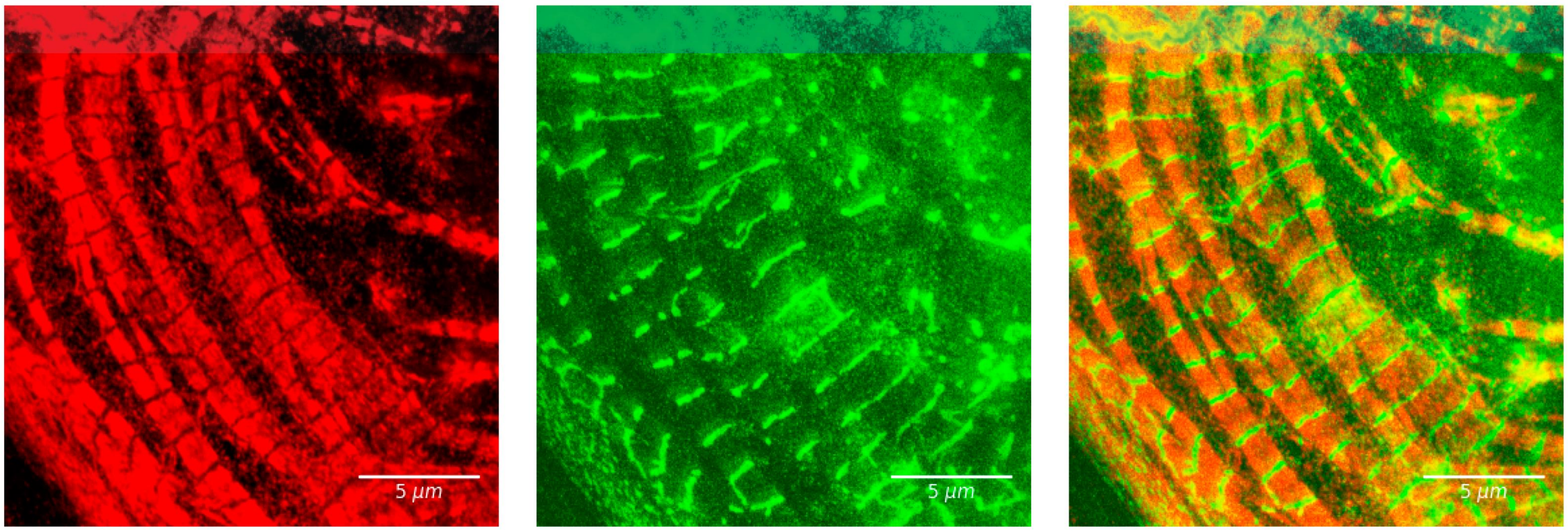
